# CDeep3M-Preview: Online segmentation using the deep neural network model zoo

**DOI:** 10.1101/2020.03.26.010660

**Authors:** Matthias G Haberl, Willy Wong, Sean Penticoff, Jihyeon Je, Matthew Madany, Adrian Borchardt, Daniela Boassa, Steven T Peltier, Mark H Ellisman

**Affiliations:** National Center for Microscopy and Imaging Research, School of Medicine, University of California San Diego, Biomedical Science Building, 9500 Gilman Drive, La Jolla

## Abstract

Sharing deep neural networks and testing the performance of trained networks typically involves a major initial commitment towards one algorithm, before knowing how the network will perform on a different dataset. Here we release a free online tool, CDeep3M-Preview, that allows end-users to rapidly test the performance of any of the pre-trained neural network models hosted on the CIL-CDeep3M modelzoo. This feature makes part of a set of complementary strategies we employ to facilitate sharing, increase reproducibility and enable quicker insights into biology. Namely we: (1) provide CDeep3M deep learning image segmentation software through cloud applications (Colab and AWS) and containerized installations (Docker and Singularity) (2) co-hosting trained deep neural networks with the relevant microscopy images and (3) providing a CDeep3M-Preview feature, enabling quick tests of trained networks on user provided test data or any of the publicly hosted large datasets. The CDeep3M-modelzoo and the cellimagelibrary.org are open for contributions of both, trained models as well as image datasets by the community and all services are free of charge.

## Main

New deep neural networks are developed rapidly and a startling number of trained models are available online for a wide range of image enhancement and analysis tasks (see ^1^ for a recent review). Since training new models is however expensive and typically requires laboriously expert annotated training data, innovations in sharing trained models effectively are critical to reduce time and cost in research. Model zoos and GitHub repositories with different networks and/or trained models are currently the most common way to share models, but are fairly disparate from the typical workflow or processing pipelines of biomedical labs. Passively hosted model zoos do not offer an immediate entry point to evaluate the performance of a neural network or a trained model, instead require to go first through complex installations - usually on high performance systems - before being able to know if the network will be useful for the specific question at hand. The computations to test a deep neural network typically require installation of several requisites on a high-performance GPU-equipped system and familiarization with the individual processing routines and configurations employed. Therefore, the use of model zoos has not been able to eliminate a major time commitment required to reproduce results. Even recent developments of more user-friendly solutions for running deep neural networks for image segmentation^2–4^ on local or cloud resources do require a time commitment of researchers at different levels to: set up, configure and troubleshoot then familiarize themselves with software settings and testing parameters and their influence on performance. Further training and testing different models on several small or individual very large imaging datasets can be cumbersome. As a result, many cutting-edge developments for image analysis with deep learning are still not used by a large portion of the biomedical imaging community.

To facilitate sharing, increase reproducibility and enable quicker insights into biology we employ a set of innovative complementary strategies: (1) we recently released a deep neural network platform, CDeep3M^4^, which circumvents installation issues and hardware requirements for end-users. We now provide a docker container of CDeep3M2 as well as a Google Colab installer with GUI (2) we are hosting CDeep3M pre-trained models in a public database (modelzoo), on cellimagelibrary.org (CIL) that also hosts relevant large microscopy datasets^5^ and (3) we are now releasing an online CDeep3M-Preview feature, offering instantaneous testing of any trained neural network that is hosted on the CIL database. This allows users to ‘test drive’ CDeep3M models within minutes on either their own data or on any region of interest on a large number of publicly hosted imaging data to determine if a trained model of interest performs well for their purpose and/or dataset (**Figure 1a-g, Supplementary Figure 1a-b**). All results are displayed through a web interface, accessible to download and can be shared with a link (**Figure 1e-g, Supplementary Figure 1c**). Users are then guided through different options how to run the same model on a larger scale or re-train the model with their own data using one of our distributions (**Figure 1h**). At the same time the CDeep3M model uploader further provides users with a way to share their trained models with the community in the modelzoo in a common place in a fully functional and testable state (**Figure 2a**).

**Figure 1.**
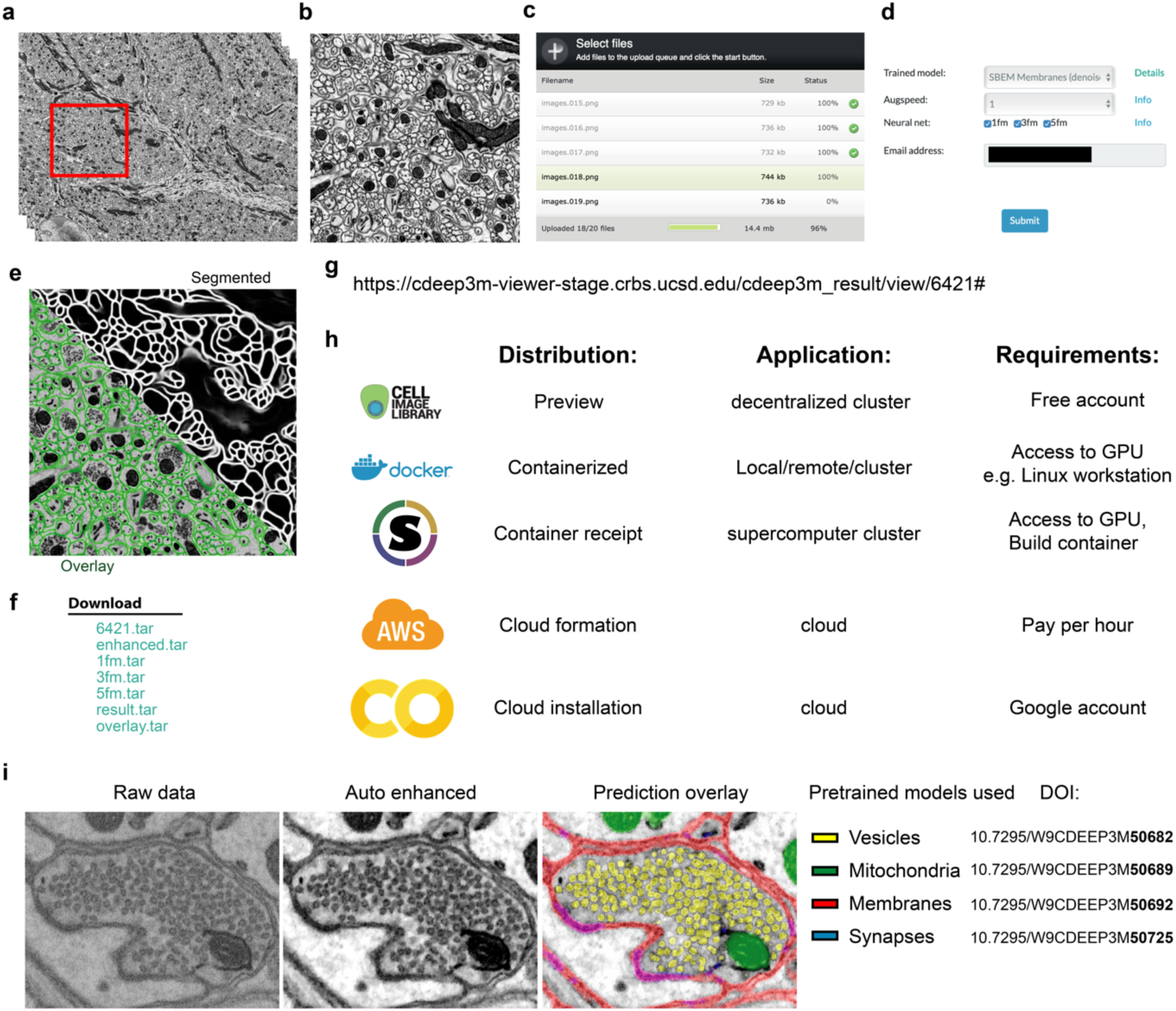
Application of CDeep3M, using the preview function. (a-b) Selection of a region of interest (ROI) from a large dataset to run CDeep3M-Preview. (c) Upload ROI data through web interface (https://cdeep3m-stage.crbs.ucsd.edu/cdeep3m). (d) Chose trained model from CDeep3M modelzoo and select parameters to perform preview. (e) Results are accessible through a web interface (here shown segmented and overlay) and (f) data is accessible for download. (g) Through the web interface the link can easily be shared with collaborators. (h) CDeep3M is available in multiple distributions to facilitate access for many groups, for quick entry points (CDeep3M-Preview, Docker, AWS, Colab), small scale tests (Preview, Colab) as well as large scale data science projects (Docker, Singularity, AWS). The end-user can then either apply the trained model to the large dataset or re-train the model with specific training data through one of those distributions. (i) Multiple pre-trained models are available on the CDeep3M-modelzoo and were combined in this example (without re-training the network for this dataset) to segment the cellular constituents of synapses. Auto-enhancement is performed in CDeep3M2 providing generalizable models.

**Figure 2.**
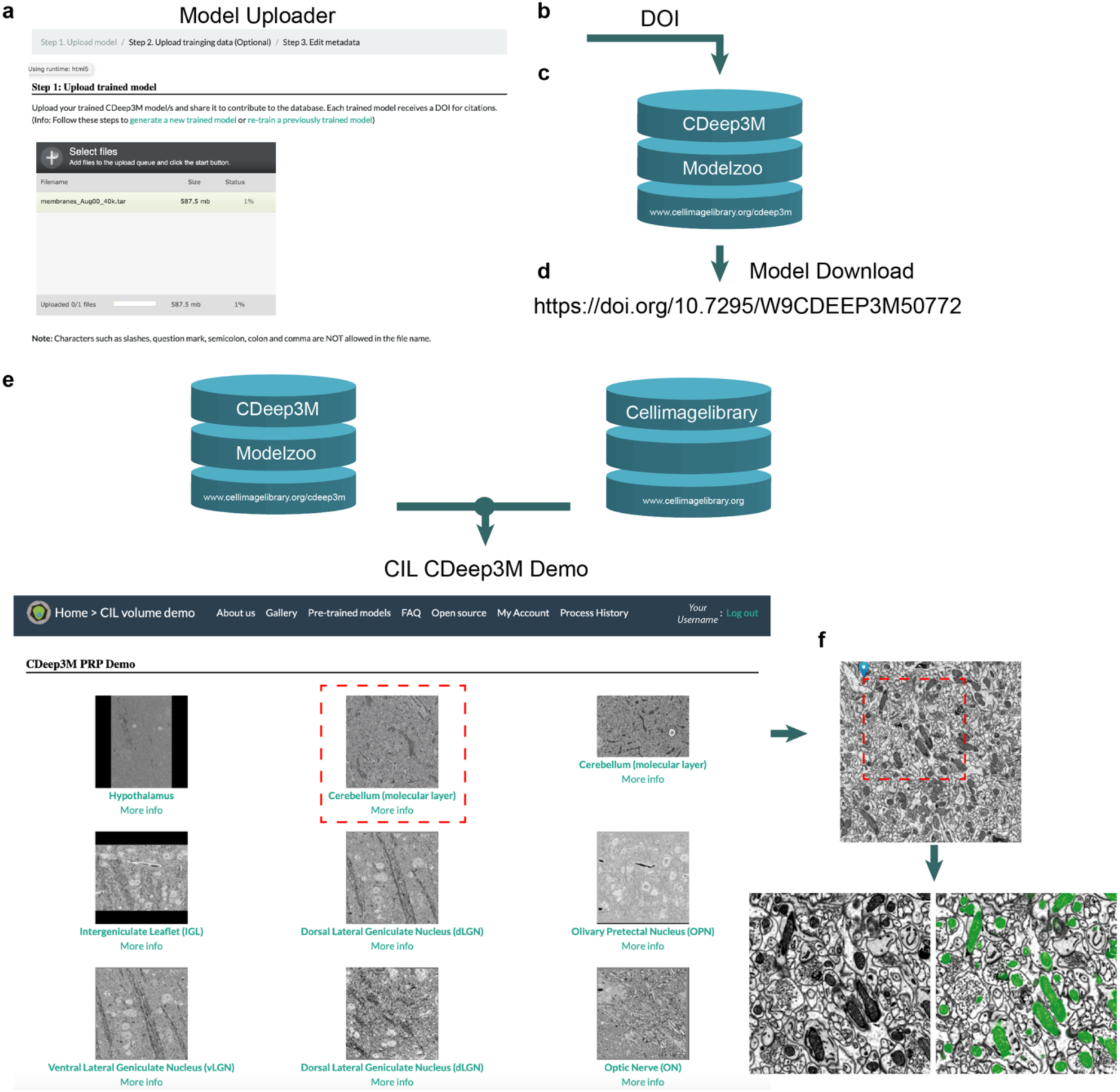
CDeep3M model uploader, database and demo function. (a) Community contributions to the CDeep3M-modelzoo are facilitated through a web interface to upload and add metainformation to trained models. (b) Each trained model receives a citable DOI and (c) is added to the CDeep3M database, and becomes therefore accessible for the Preview function and (d) can be easily downloaded. (e) The cellimagelibrary hosts many large-scale imaging datasets to which any of the trained networks in the modelzoo can now be applied through the CDeep3M-demo function in an image browser. (f) Example using a broadly trained CDeep3M model on a dataset for which it was no trained (before any transfer learning is applied). Results can be easily shared through using the specific job ID and the web interface. Results from (f) are at: https://cdeep3m-viewer.crbs.ucsd.edu/cdeep3m_result/view/6447

Data processing with state-of-the-art deep neural networks requires high-end hardware with GPUs with sufficient vRAM. Online processing using deep neural networks for many users, as provided by CDeep3M-Preview, is limited by hardware availability equipped with high-end GPUs. We therefore implemented a scheduling system outsourcing the processing of the preview to the infrastructure of the Pacific Research Pipeline (PRP), a decentralized computing cluster with GPUs and storage nodes. The PRP cluster is managed by Kubernetes, using containerized applications to rapidly deploy computing jobs to available hardware. Our installation for the CDeep3M-Preview is containerized as a Docker image on the PRP cluster, and is streamed to available nodes allowing for near instantaneous start-up times (within seconds). The imaging data and pre-trained models are sent from CIL and the commands to initiate segmentation and the subsequent overlay with the segmentation is submitted. Using a next-generation decentralized computer cluster, rather than running the backend processing on workstation/s, provides scalability at times of high demand. In practice, this means that the end users can test whether a trained model performs on their dataset in less than 5min total, without programming knowledge and without any hardware requirements (**Figure 1**).

Once the user has identified a pretrained CDeep3M model and tested settings which perform well for a dataset of interest (**Figure 1e**), several routes are made available that provide quick implementations for applying the pre-trained network and the settings on the complete large-scale dataset. To this end we maintain several preconfigured installers for the use on local hardware, on supercomputer clusters or on cloud providers. Following pre-configured CDeep3M installers are provided: (i) Docker container (ii) Singularity recipe (iii) AWS cloudformation template (iv) Colab Notebook (**Figure 1h**). Detailed descriptions for configurations of each of those solutions are available with the links provided. Without much effort or the obstacle of requiring their own GPUs or funding for high-performance hardware the users can now also take advantage of the free GPUs provided Google Colab, with the CDeep3M- colab installer and graphical user interface (**Supplementary Figure 2**).

It is advantageous to co-host trained models and imaging data on the same platform and maximize cross-linking between the two and facilitate testing across several large datasets for generalizability. With the new extensions to the CDeep3M-Modelzoo the users can upload their trained models to the CIL repository (**Figure 2a-d**), in order to share them or to apply them through the preview function on one of the imaging datasets. In addition, metadata about the available trained models is stored in the database, such as the targeted cell component, staining procedures, imaging modality and voxel dimensions. When releasing a trained model on the CIL database to the public, it will generate a citable Digital Object Identifier (DOI) (**Figure 2b, 2d**). The DOI is a persistent identifier used to identify objects uniquely, standardized by the International Organization for Standardization (ISO). To help other groups unlock valuable large scale data we are providing the CDeep3M-Demo, facilitating to test pre-trained models on areas of interest on the large scale datasets available on http://cellimagelibrary.org/ (CIL; **Figure 2e**). We co-host trained CDeep3M models on the CIL providing us with the infrastructure already in place for data storage, metadata organization and large-scale image visualization. CIL is open-source software providing storage and user interfaces to deliver a publicly searchable database of microscopy images and metadata to facilitate data sharing and reuse. The CIL data input form allows end-users to submit images to the CIL data repository and annotate the images with the ontology markup.

Together with the online preview function we release an upgrade to CDeep3M2, which provides additional functionalities and reduced runtimes. The new version of CDeep3M is backwards compatible, so that all previously trained models can still be applied and used for transfer learning with the new release. Importantly, we incorporated enhanced image augmentation strategies, that can easily be configured, to facilitate the training of more broadly tuned models. In the enhanced training augmentation pipeline, the images are first processed through the sixteen rotations and inversions (x/y and z) before each stack will go through a set of additional secondary and/or tertiary augmentations. The augmentations are performed as follows: primary augmentations consisting of rotations and inversions (x: left/right, y: top/down, z: forward/reverse) are always performed, secondary augmentations consisting of image filters (contrasting, sharpening, blurring, total variation denoising, introduction of uniform noise, histogram equalization, skewing, elastic distortion; and tertiary augmentations, resizing the images. Secondary and tertiary augmentation strengths are determined by the user with a scaling factor between 0 (no additional augmentation) and 10 (strong augmentation). The combination of augmentation strength can be customized for each dataset depending on the purpose of the training (fine tuning of model for one individual dataset or broad training for generalizable model). Furthermore, users can now easily provide multiple training datasets that will be used during training, which facilitates generating broadly applicable trained networks.

Applying the CDeep3M-Preview and Demo functions on the cellimagelibrary serves us as an extreme test case scenario at unprecedented scale to test and improve how well a trained deep neural network performs on previously unseen data (generalizability). The available datasets are stored at various imaging conditions (pixel- and voxelsize), ranging from 8bit unsigned integer to 32bit signed integer, with different levels of noise, staining intensities and imaging conditions, acquired from different tissue types. These constraints are very typical for different biological sample preparation and imaging. Rather than performing training for each individual dataset, we focused on training and providing more generalizing models, and stabilize their performance through improved image pre-processing, which will reduce the requirements to re-train the models. Mainly we automatized the following steps: image conversion to 8bit, with simultaneous clipping of outlier pixels, readjustment of contrast and a total variation denoising (**Figure 1i**). Overall, we note that these implementations improved the generalizability of trained models making them broadly applicable to many more datasets, since this reduces most of the extreme variations in signal-to-noise levels and contrast between different SBEM datasets.

Altogether, the CDeep3M-Preview and new set of tools provided here gives biomedical researchers immediate access to experiment with AI for image segmentation and the ability to test different trained models near-instantaneously. A similar quick entry approach has been taken recently with DeepCell, which allows users to track cells in their own live cell imaging data with deep learning^6^. On the CDeep3M modelzoo a broad range of pre-trained models, trained on segmentation tasks for electron microscopy, X-ray microCT and light microscopy data are readily available. Furthermore, by taking advantage of an emerging cyberinfrastructure of decentralized compute cluster, the preview is scalable to account for the high demand of many users. This approach can serve as an entry point for community members with no experience in deep or machine learning to become familiar with the technology and experiment with the effect of different parameter settings and will contribute to democratize deep learning in the bioimaging community while allowing them to scale their use case afterwards quickly to extremely large datasets through the CDeep3M built in processing pipelines.

## Acknowledgements

This research was funded by grants from the National Institutes of Health under award numbers 3R01GM082949, 3P41GM103412, 1RF1MH120685, 1R01AG065549 and 5R01DA038896. The CDeep3M-Preview and Demo are using PRP/Chase-CI, which is supported by its members institutions and the United States National Science Foundation through the NSF awards CNS-1456638, CNS-1730158, ACI-1540112, ACI-1541349, and OAC-1826967.

## Author contributions

M.G.H. performed computational experiments. M.G.H. and J.J. wrote CDeep3M upgrade. W.W. implemented CDeep3M-Preview and CDeep3M-Demo interface on CIL and model zoo uploader and database. S.P. implemented preview backend scheduler. A.B. maintains singularity recipe and AWS cloud formation template. M.G.H. wrote colab implementation with GUI. M.G.H. and M.M trained models hosted on modelzoo. D.B. acquired SBEM data. M.G.H., S.T.P. and M.H.E. supervised project. M.G.H. wrote manuscript with input from all authors.

## Competing interests

The authors declare no competing interests.

## Methods

### CIL backend system

Under the CIL system, the metadata will be converted into JSON format and stored in a NoSQL datastore. The CIL utilizes Elasticsearch as its JSON search engine since it provides a distributed, multitenant-capable full-text search engine with an HTTP web interface. When JSON data is imported into the Elasticsearch datastore, all of the data fields are automatically indexed and immediately searchable using the built-in web-service. These built-in functions in Elasticsearch are crucial for software development because it saves tremendous development time otherwise spent building data models and backend services.

### Backend operations Scheduling system

At the core of the scheduling system for the CDeep3M-Preview and Demo functions is the beanstalkd queue (https://beanstalkd.github.io/). Beanstalkd is a robust multichannel FIFO queuing system. A single queue (or tube) contains the jobs in the order they are submitted. Each job consists of the UID of the requestor, a description of the job to be run and its arguments, and an authentication token with a timestamp. A job can be in one of 4 states, ready, reserved, delayed, or buried. A costum written Perl based web API (stalker_web) is used as an abstraction layer for the interaction with beanstalkd. A job is submitted to the system via a client; the client does a few basic sanity checks against the submitted job, secures a token and submits the job to the queue via stalker_web and if the submission is successful is returned the job id in the tube. Once in the tube the job is in the ready state.

On the processing side there is a worker that periodically checks the tube for ready jobs, if a job is found, it’s parsed and if the worker has the capabilities needed to run the job it verifies the token then reserves the job and then places it in the delayed state for an amount of time equal to the expected runtime (ERT). This removes the job from the ready state so no other workers will see the job. Since these are typically longer running jobs and are designed to run on geographically disparate hardware in an ad libitum fashion the delay is set to remove the need for the constant communication between worker and queue required by the reserved state.

The worker then sets the environment, downloads any data or models it needs, and proceeds to process the job as the UID of the submitter, collecting the output of the commands into a log. A separate watchdog process is started that will check every 90% of ERT if the worker process is still running and if it is, try to reset the delay on that job to ERT until successful or the parent process exits. Once the commands finish, the worker deletes the job from the queue and the log is published to a separate tube named with the job id of the original job.

The worker sends the output data back to the requestor via an API call. If the worker should die or communication between the worker and the queue goes down the delay time will eventually run out and the job will go back into ready state for another worker to pick up and process.

### PRP platform

CDeep3M-preview is running as a Kubernetes Pod on the Pacific Research Platform (PRP) and configured to use gpu-pods equipped with GPUs with least 11GB vRAM. PRP is spanning over 20 universities and institutions, all connected by dedicated optical light-paths at speeds of 10-100gb/s. A list of currently available PRP resources can be found at: https://ucsd-prp.gitlab.io/userdocs/running/gpu-pods/.

### CDeep3M2 data augmentations

Data augmentations are used to avoid overfitting to training data and intended to increase generalizability of trained models. To facilitate regulating which data augmentation strategies are used we chose to separate augmentation strategies into following three categories:

*Primary augmentations* refer to augmentations that only change the image orientations, such as data rotations and flipping in x, y and z directions. Since those leave the data unaltered they are always performed, on each training dataset, to generate 16 variations of the same data.

*Secondary augmentations* are data augmentations which alter the noise level, the brightness or the contrast of the images. Secondary augmentations are performed if a setting between 1 (weak) to 10 (strong) is chosen. Following operations are used in secondary augmentations: increase and lowering of the image contrast; sharpening and gaussian blurring of the image stack; total variation using the Chambolle and/or Bregman filter to denoise the image; adding of random binary gaussian noise; normalization of the image by performing histogram equalization; random skewing of the image stack to four different directions (upper left, upper right, lower left, lower right) while maintaining the image’s aspect ratio; elastic distortions across the image stack in which the first and the last images of the stack are distorted by a randomly generated gaussian vector field while the images in between are distorted by the interpolated field in between the two.

*Tertiary Augmentation* are performed if a setting between 1 (weak) to 10 (strong) is chosen. During the tertiary augmentations images are resized, according to a strength selected by the end user (values 1-10). Data is resized using upscaling alternating with downscaling, to broaden the networks capabilities to recognize the same object at a different pixelsize.

## Code availability

All code is open access. The CDeep3M2 Docker container can be pulled directly from Docker-Hub at https://hub.docker.com/r/ncmir/cdeep3m or simply running ‘docker pull ncmir/cdeep3m:latest’. The Google Colab Jupyter Notebooks with graphical user-interface (GUI) are available on GitHub at https://github.com/haberlmatt/cdeep3m-colab. The CDeep3M2 AWS cloudformation template is available <u>here</u>. CDeep3M2 source code is available on GitHub https://github.com/CRBS/cdeep3m2. The singularity image is available at: http://cellimagelibrary.org/cdeep3m/singularity.

