## Supplementary Figures for "CDeep3M-Preview: Online segmentation using the deep neural network model zoo"

### Supplementary Material

#### Supplementary Figure 1

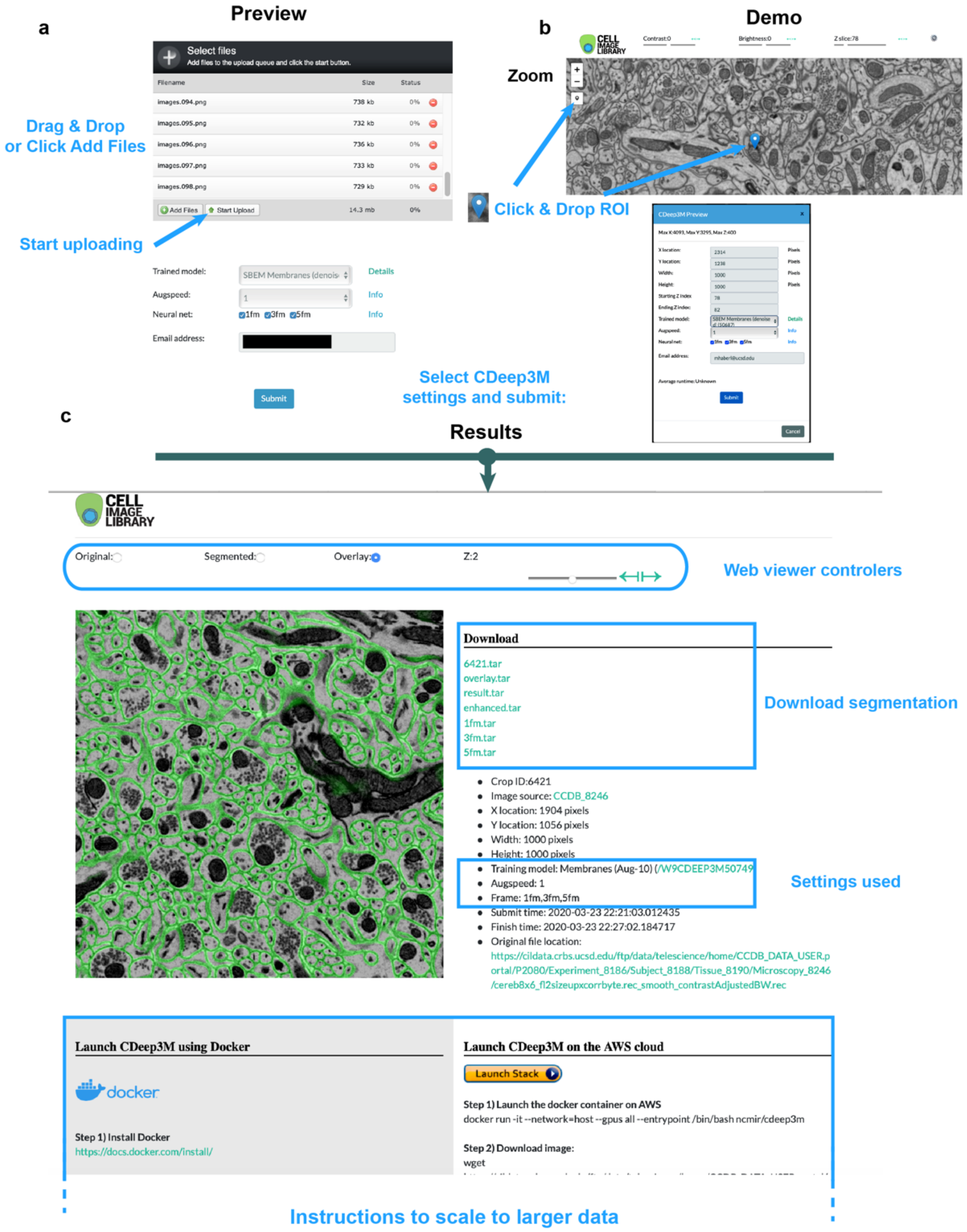


**Supplementary Figure 1. CDeep3M online segmentation.** (a) CDeep3M-Preview allows end-users to submit their own images to test the performance of any trained model from the CDeep3M modelzoo. (b) CDeep3M-Demo allows the users to perform segmentation tasks on large-scale data stored on the cellimagelibrary.org using trained CDeep3M models. (c) The results of CDeep3M-Preview and Demo are accessible through a web image browser, can be shared and downloaded. Instructions to perform the segmentation as done in those tests on large scale are provided. The online segmentations are free of charge to the end-user.

#### Supplementary Figure 2

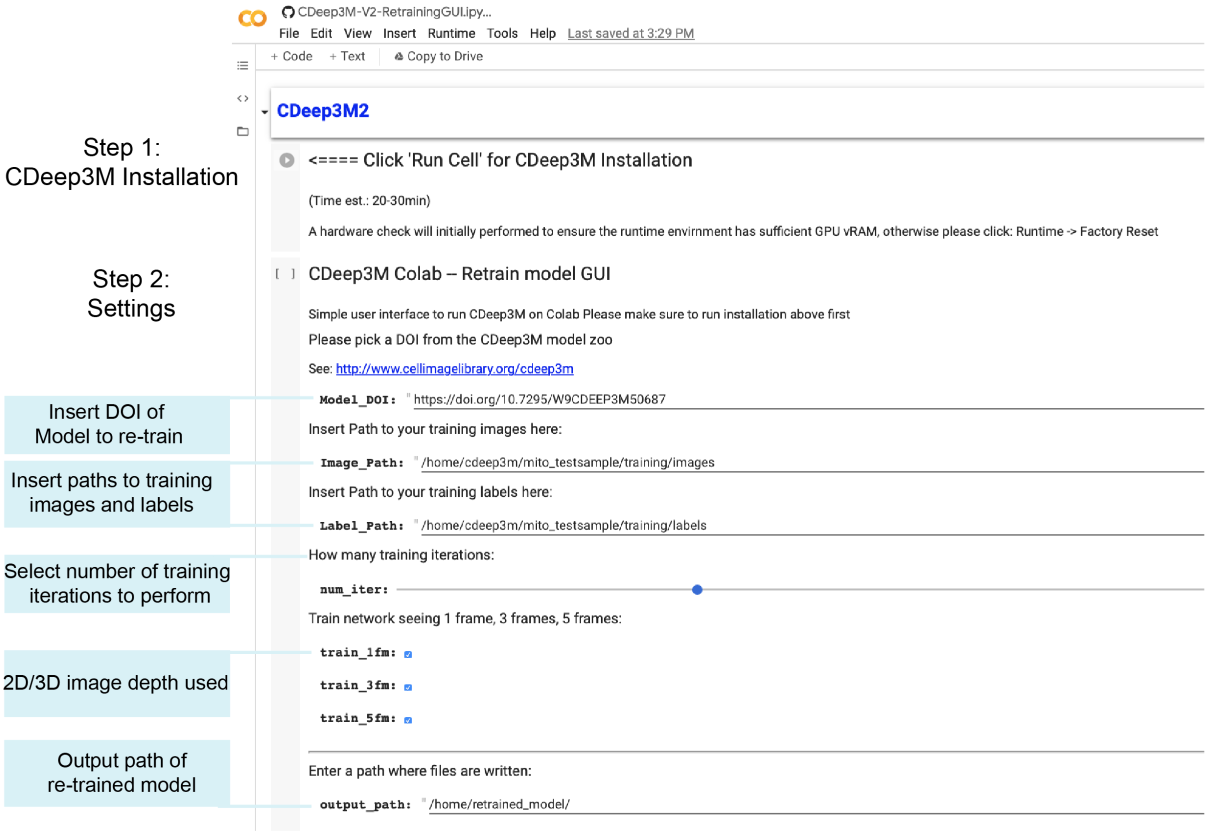


**Supplementary Figure 2. CDeep3M-Colab GUI for re-training a model.** Transfer learning can be applied to a previously trained CDeep3M model for free using the provided notebook. In the first step all requirements of CDeep3M. In the second step several parameters are inserted by the user, such as the model on which transfer learning should be applied, the location of the new training images and the settings of the training that will be performed. Additional notebooks are available to perform all other CDeep3M capabilities. For clarity the GUIs to perform different tasks are separated into different notebooks. All functions are also accessible through the command-line-interface on CDeep3M-Colab. For more info see: https://github.com/haberlmatt/cdeep3m-colab
